# LINC00536 regulates transcriptional repressor TRPS1 in breast cancer

**DOI:** 10.64898/2026.08.31.748275

**Authors:** Rui Zhang, Jiarong Li, Dunarel Badescu, Jiannis Ragoussis, Richard Kremer

## Abstract

Metastatic breast cancer with complex molecular mechanisms of progression accounts for most cancer related deaths in women. To improve diagnosis and drug development, it is important to identify novel biomarkers and critical molecular pathways involved in tumor initiation and progression. Here, we profiled and analyzed the expression of long non-coding RNAs (lncRNAs) from three distinct stages of tumor initiation and progression (hyperplasia, adenoma, and carcinoma). We performed RNAseq on tumor and mammary epithelial cells derived from ROSAmT/mG tumor and non-tumor mice. We identified 1913 differentially expressed protein coding genes and 324 lncRNAs in breast cancer cells of all stages compared with normal mammary epithelial cells. Pearson correlation analysis correlated 93 differentially expressed lncRNAs with protein coding genes, providing a comprehensive lncRNA-protein coding genes co-expression network. Among them, we focused on *Gm19303* which was paired with the differentially expressed protein coding gene, transcriptional repressor GATA binding 1 (*Trps1*), and identified its human counterpart as *LINC00536*. Both *LINC00536* and *TRPS1* are only overexpressed in breast cancer and correlate with poor prognosis of patient from the TCGA and GTEx databases. Single cell RNAseq data from the Atlas of Human breast cancers further confirmed that TRPS1 is upregulated in human breast cancer compared to normal human mammary tissue with highest expression in ER^+^ subgroup. In summary, our study explored the potential role of lncRNAs in breast cancer initiation and progression.

**Implications statement:** Our findings imply that human LINC00536/TRPS1 serves as a novel and early biomarker of cancer progression and a potential therapeutic target for breast cancer.

## 1. Introduction

Breast cancer (BC) is the most frequently diagnosed cancer and the leading cause of cancer-related mortality among women of all ages ^1^. Based on the well-defined histological types and protein markers, BC can be defined into three clinical subtypes: hormone-receptor (HR)-positive (HR^+^; ER^+^, PR^+/-^ and HER2^−^), HER2-positive (HER2^+^), and triple-negative (TN; ER^−^, PR^−^ and HER2^−^) ^2^. According to the PAM50 gene signature ^3^, BC is also stratified into five molecular subtypes: luminal-like (LumA and LumB), HER2-enriched (HER2E), basal-like, and normal-like. Luminal A represents around 50% to 60% of all BC cases and is frequently associated with better prognosis, in contrast to luminal B or non-luminal subtypes ^4^. While PAM50 has provided important insights into prognosis and treatment, the molecular mechanisms involved in therapeutic resistance and recurrence have not yet been fully understood.

Genetically engineered mouse models (GEMMs) have been widely used to study the molecular events involved in BC initiation, progression, and metastasis. Polyoma middle T-antigen (PyMT) mouse model remains the commonly used GEMM due to the rapid development of multifocal tumors and lung metastasis in all the animals. First described by Muller in 1992, the expression of PyMT from the mouse polyoma virus is under the control of the mouse mammary tumor virus’s long terminal repeats promoter (MMTV-LTR) and is restricted to mammary epithelium ^5^. Despite not being a human oncogene, PyMT mimics receptor tyrosine kinase signaling which are commonly activated in many human malignancies including BC ^6^. The primary tumors developed in this mouse model progress through four stereotypical stages of cancer progression that mimic human ductal BC progression - hyperplasia, adenoma/mammary intraepithelial neoplasia (MIN), early and late carcinoma – from premalignancy to malignancy ^7^.

In recent years, lncRNAs emerged as a novel class of regulatory molecules in cancer ^8^. LncRNAs belong to a large class of noncoding RNAs with a typical length of more than 200 nucleotides. The most updated annotation (GENCODE 46) indicates that there are more than 17,900 lncRNA genes and approximately 48,000 lncRNA transcripts in the human genome, with more and more novel lncRNAs being discovered ^9^. The biogenesis of lncRNAs sometimes has similar structures to that of protein coding genes (PCGs) such as 5’ modified caps, exons and poly-adenylated tails at their 3’-end ^10^. LncRNAs are spliced post-transcriptionally, but lack functional open reading frames, and are considered incapable of encoding functional proteins ^10^. *H19* is one of the most studied lncRNAs involved in BC progression, which is aberrantly upregulated in human breast tumor tissues and cells and is associated with an increased risk of BC ^11^. LncRNA *HOTAIR* is aberrantly expressed in BC compared to normal mammary epithelium ^12^. Higher level of *HOTAIR* expression is correlated with a poor prognosis for BC and is an independent biomarker for predicting BC mortality and metastasis ^12^. *MALAT1* was first found to be associated with progression in patients with squamous cell cancer and adenocarcinoma of the lung ^13^. Wu *et al.* found that an elevated expression of *MALAT1* in BC compared to non-cancer breast tissues and has its highest expression in metastatic TNBC and trastuzumab-resistant HER2 overexpressing cells ^14^. As only a handful of the almost 20,000 annotated lncRNAs (GENCODE 46) have been characterized and the temporal expression profile of lncRNAs in BC initiation and progression is still unclear, it is essential to perform an unbiased RNA sequencing (RNAseq) screen to identify the lncRNAs that exhibit aberrant expression in mammary tumors.

In this study, we aimed to examine the expression of lncRNAs in the full range of breast tumor development. The availability of the conditional fluorescent reporter mouse model (MMTV-Cre/mT/mG) has enabled us to trace Cre expression exclusively in mammary epithelial cells ^15^. Here, we combined the classic PyMT-induced BC mouse model with MMTV-Cre/mT/mG, which allowed us to specifically trace and enrich the GFP^+^ mammary epithelial cells from the early stage of tumor development to late-stage carcinoma. We performed RNAseq analysis and achieved a comprehensive and unbiased gene expression comparison between primary mammary tumor and normal mammary gland epithelial cells at three distinct stages of BC progression. We carried out differential gene expression profiling and time-course analysis to identify novel candidate lncRNAs that may be linked to BC initiation and progression. Thus, we propose that these temporally aberrantly expressed lncRNAs might act as potential biomarkers and therapeutic targets for BC.

## 2. Materials and Methods

### 2.1 Mouse Lines, Breeding, and Genotyping

All animals were housed in the Research Institute of the McGill University Health Centre (MUHC) Animal Resources Division, and all animal protocols were reviewed and approved by the Glen Facility Animal Care Committee (FACC). MMTV-PyMT ^5^ and MMTV-Cre mice ^16^ (RRID: IMSR_JAX:003553) on a pure FVB/NJ (RRID: IMSR_JAX:001800) background were kindly supplied by Dr. William Muller (McGill University Cancer Center). ROSA mT/mG reporter mouse line (mT/mG) was obtained from the Jackson Laboratory (Bar Harbor, Me, RRID: IMSR_JAX:007576). The ROSA mT/mG reporter mouse on C57BL/6J background was backcrossed onto the FVB/NJ background (ten generations) and resulting strain was 99 percent FVB/NJ. For lineage-tracing and RNAseq experiments, MMTV-PyMT; MMTV-Cre^+^ mice or MMTV-Cre^+^ non-cancer mice were crossed with ROSA mT/mG mice. The genotyping of the PyMT and Cre transgene were performed by standard PCR using primer sets as described previously ^17^. Genotyping was performed on DNA isolated using standard protocols from tail snips obtained at or just before weaning of litters. In brief, PCR reaction was carried out as follows: 94 ◦C for 30 s, 64 ◦C for 1 min, 72 ◦C for 1 min, 35 cycles. The IVIS® Spectrum *in vivo* imaging system was applied to measure the tail epifluorescence which is sufficient to identify ROSA mT/mG mice from wildtype mice. Mice were tested under three stages with three biological replicates per stage. Comparisons under these conditions are explained in Figure 1B.

**Figure 1.**
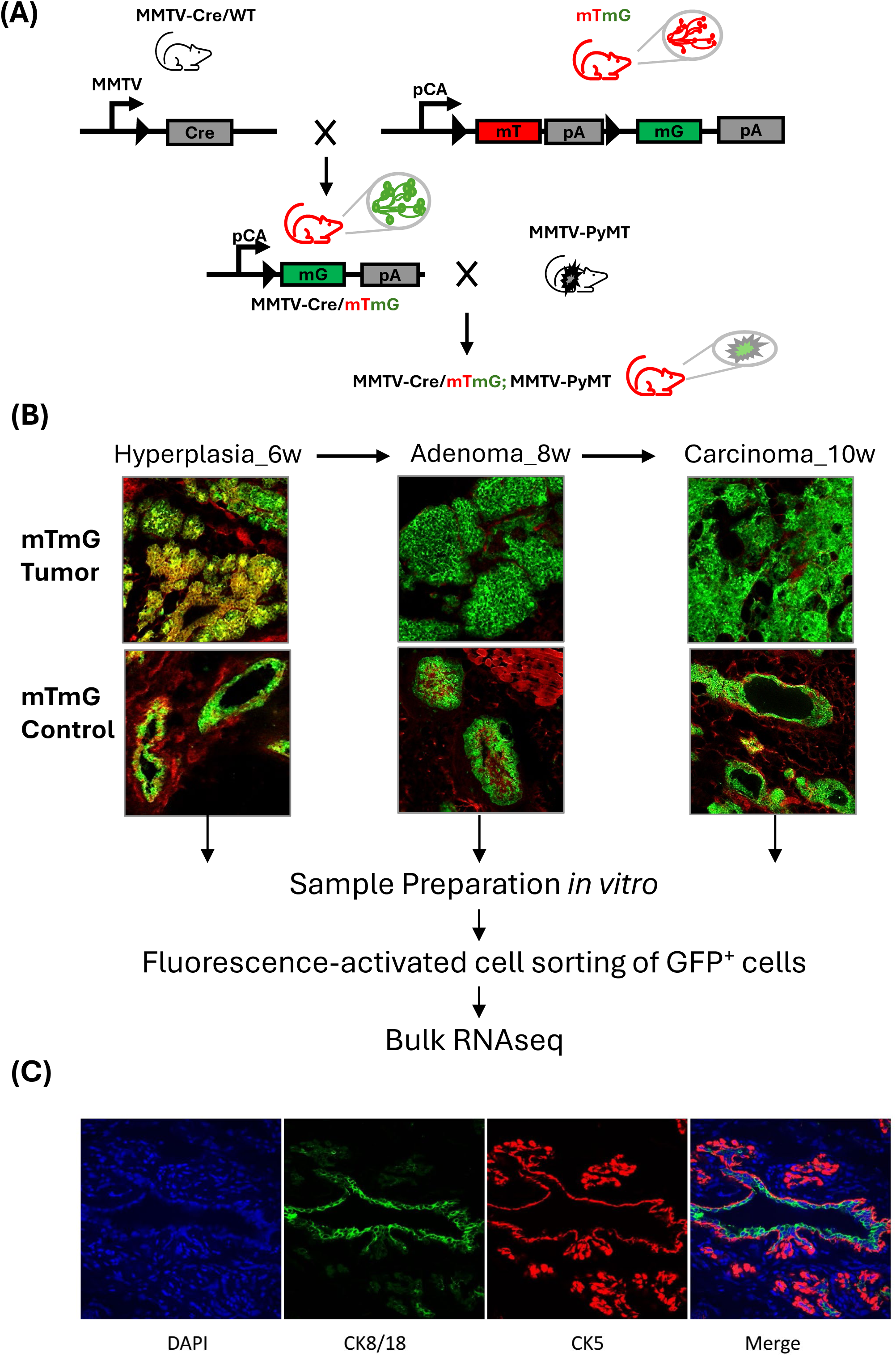
Generation mT/mG control and mT/mG tumor Mouse Model. **(A)** Schematic diagram of the genetic construction of mT/mG control and mT/mG tumor mice. **(B)** Ubiquitous expression of mTd and tissue-specific expression of GFP in ducts and alveoli in the mammary gland from mT/mG control mouse and expression of GFP in breast cancer cells from mT/mG tumor mouse (fresh tissue). A diagram illustrates the workflow for RNAseq including sample preparation, GFP^+^ cell sorting and RNA extraction. **(C)** IF staining of mammary tissues from 8-week-old MMTV-PyMT mice shows that CK8/18 is specifically expressed in epithelial cells (green), while CK5 is expressed in basal/myoepithelial cells (red), which form the outer layer of the ducts and lobules regions. Scale bar: 20X.

### 2.2 Confocal Microscopy, Immunofluorescence (IF), and Immunohistochemistry

Fresh tissues were dissected from ROSA mT/mG; MMTV-Cre⁺ tumor and non-tumor mice. Tissues were embedded in Optimal Cutting Temperature (OCT) compound, placed into OCT molds, and frozen in a dry ice container. Tissue sections (10 µm thick) were cut using a cryostat (Histopathology Platform in Research Institute of MUHC). The cryosections were stored at −80 °C for several hours, then thawed at room temperature for 10-20 minutes. Slides were rehydrated in wash buffer for 10 minutes and subsequently processed using a standard IF staining protocol ^18^.

IF staining was performed according to the manufacturer’s protocol (Invitrogen, ON). Briefly, after antigen retrieval, tissue sections were blocked with 10% serum in PBS for 30 minutes, washed for 3 minutes in PBS, and incubated overnight at 4 °C with primary antibodies: rabbit anti-human TRPS1 (Thermo Fisher, PA5-49890), guinea pig polyclonal anti-Keratin K8/K18 (Progen, GP11), mouse monoclonal anti-Ki67 (Cell signaling Technology, #9449), and mouse monoclonal anti-CK5 (Abcam, ab52635). After washing with PBS (5 minutes), slides were incubated for 45 minutes in the dark with the appropriate secondary antibodies: goat anti-rabbit Alexa Fluor 546, goat anti-mouse Alexa Fluor 568 (A-11004, Invitrogen, Burlington, ON), and sheep anti-guinea pig Alexa Fluor 488. Nuclei were counterstained with 4ʹ,6-diamidino-2-phenylindole (DAPI, Invitrogen, ON) for 10 minutes in the dark. Slides were washed three times in PBS and mounted with aqueous mounting medium (Thermo Electron, Pittsburgh, PA). Images were acquired using a Zeiss LSM780-NLO Laser Scanning Confocal with IR-OPO lasers microscope, a 10X/0.30 WD=5.2 objective, and Zen2012 image acquisition software (Molecular Imaging Platform in Research Institute of MUHC).

### 2.3 Isolation of Primary Mammary Gland and Breast Tumor

Primary tissues (tumor or gland) were isolated from mammary by stepwise mechanical disruption and enzymatic digestion according to our published protocols ^17^. Tissues (tumor or gland) were harvested from 6- to 10-wk-old mice, minced with a scalpel, and incubated for digestion for 2 h at 37 °C with gentle rocking in FBS free DMEM (Multicell, Wisent Inc., St. Bruno, Quebec, Canada) supplemented with 2.4 mg/ml collagenase B and 5U/ml Dispase II (4942078001, Roche, Switzerland). Tissue fragments were washed with PBS, centrifuged, and resuspended in DMEM with 10 % Fetal Bovine Serum (Gibco, US Origin), 100 IU/mL penicillin, and 100 ug/mL streptomycin (Multicell, Wisent Inc., St. Bruno, Quebec, Canada).

### 2.4 Fluorescence-Activated Cell Sorting (FACS) of Single GFP^+^ Cells

Breast tumor cells (from MMTV-PyMT; Cre^+^; mT/mG mouse) or mammary epithelial cells (from Cre^+^; mT/mG mouse) were harvested after 48h incubation in DMEM with 10 % Fetal Bovine Serum. The resulting cell were washed in PBS, filtered through 40-μm cell strainers, and resuspended in complete DMEM prior to sorting. Following doublet exclusion using FSC-A and FSC-H scatter plot, GFP and/or tdTomato (mTd) expressing cells were sorted on a BD FACS Aria Fusion (BD Bioscience) at Immunophenotyping Platform in the Research Institute of MUHC. For the RNAseq experiments, a minimum of 20,000 single cells were sorted.

### 2.5 RNA Isolation and Sequencing

Total RNA was isolated either directly from the sorted GFP^+^ tumor cells or from mammary epithelial cells after FACS using the miRNeasy mini kit (Qiagen) according to the manufacturer’s protocol. RNA quantity was assessed by Qubit 4 Fluorometer (Invitrogen), and RNA quality was evaluated by Agilent TapeStation Laptop (RRID:SCR_019547). For high-throughput sequencing, RNA samples were required to have an RNA integrity number (RIN) ≥ 7. The workflow from sampling to sequencing was performed according to the previously published detailed protocol ^19^ (DOI: 10.3390/cancers15153763). In summary, the RNAseq transcriptome strand library was prepared by following the TruSeqTM Stranded Total RNA kit from Illumina (San Diego, CA, USA) using 100 ng of total RNA for each sample. Briefly, ribosomal RNA (rRNA) depletion was achieved using an Illumina Ribo-Zero Magnetic kit. First strand cDNA synthesis used Super strand synthesis Act D kit containing random primers and Super Script II reverse transcriptase to convert the original mRNA to complementary cDNA. Second strand cDNA synthesis involves removal of the original RNA template and synthesizes a replacement strand, incorporating dUTP in place of dTTP to generate ds cDNA. After cDNA synthesis, a single “A” nucleotide is added to the 3’ ends of the blunt fragments to prevent them from ligating to one another during the adapter ligation reaction. During adapter ligation process, each sample had a different RNA adapter index for the identification in subsequent data analysis (Illumina® Nextera™ DNA Unique Dual Indexes). AMPure XP beads were used to purify the ds cDNA from the second strand reaction mix leaving exclusively the blunt-ended cDNA for the subsequent DNA enrichment. The PCR was performed with a PCR primer cocktail that anneals to the ends of the adapters to exclusively enrich and amplify those DNA fragments that have adapter molecules on both ends. Only the first strand cDNAs were amplified in this step. After quantification by LightCycler® 96 Instrument with KAPA library quantification kit, the paired-end RNAseq library was sequenced with Illumina NovaSeq6000 at McGill Genome Center according to the standard protocol.

### 2.6 RNAseq Analysis

The pipeline for sequencing alignment and downstream differentially expressed genes (DEGs) analysis was performed according to the previously published protocols ^19^ (DOI: 10.3390/cancers15153763). In summary, sequences were aligned using Hisat first against the mouse transcriptome as defined by Gencode gene models M16 ^20^, with default parameters and the remaining unmapped genes to the Ensembl GRCm38 reference mouse genome ^21^. Aligned reads from multiple read groups belonging to the same sample were indexed, sorted, and merged using sambamba v0.5.4 ^22^, a faster implementation of the Samtools algorithms ^23^ (RRID:SCR_002105). Amplification duplicates were removed using Picard tools v1.128 (RRID:SCR_006525). Various quality controls from the RNASEQC package were used, including the genes detected, mapping rates, duplication rates, and intronic rate, based on metrics collected for each sample used ^24^. HTSeq Count ^25^ was applied to enumerate the counts for each gene using the Gencode M16 GTF. All statistical analysis were carried out in R v3.6.2 ^26^. We performed minimal pre-filtering (keep only rows that have at least 10 reads total) to reduce the memory size of the data and increase the speed of the transformation and testing within DESeq2 ^27^ (RRID:SCR_015687). The principal components analysis (PCA) included in DESeq2. The R package DeSeq2 ^27^ was also used to perform statistical analysis on gene counts and to detect DEGs.

### 2.7 Time-Course Data Analysis

A complete list of DEGs, both upregulated and downregulated, in the tumor mice compared to non-tumor mice is presented in Figure 2. There, the union of 324 lncRNAs and 1,913 protein-coding RNAs (mRNAs) are differentially expressed in hyperplasia, adenoma and carcinoma compared with normal mammary gland at the same timepoints. To group gene expression into clusters base on similarity in their expression patterns along tumor initiation and progression, we used a K-means clustering approach. We scaled the DEGs expression and centered them to fit into the range and subjected them to the K-means clustering procedure. We plotted the number of clusters against the total within cluster SS (sums of squares) and K-means= 4 was selected as optimal. The results were visualized as line graphs using “ggplot2” library. Genes belonging to individual clusters were subjected to functional annotation analysis using “clusterProfiler” package to perform KEGG pathways enrichment analysis ^28^.

**Figure 2.**
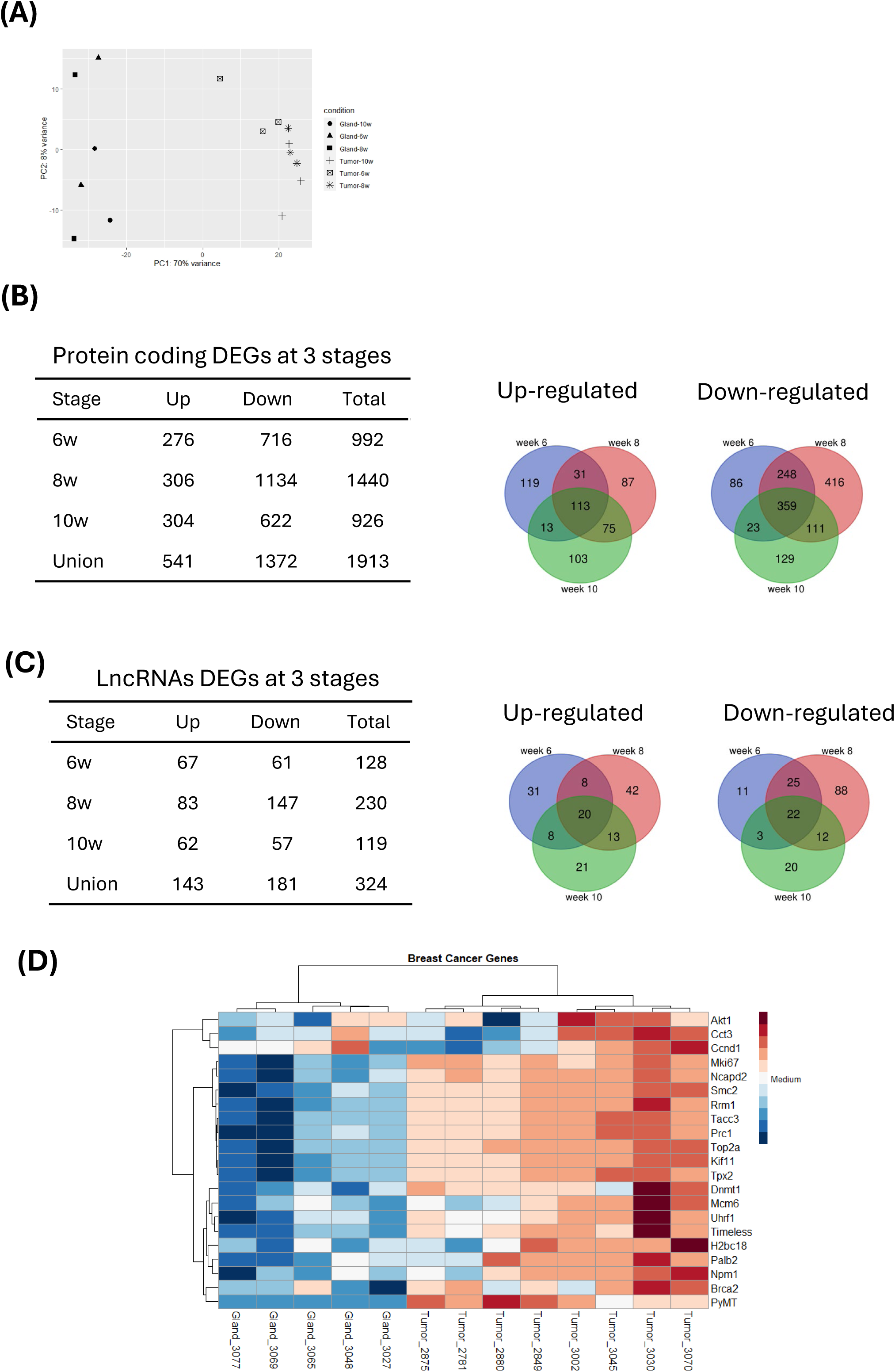
DEGs along tumor progression. **(A)** PCA of mT/mG tumor and mT/mG control samples. It shows the separation between tumor cells and normal mammary epithelial cells. **(B)** Differentially expressed PCGs and Venn diagram of differentially expressed PCGs at three stages. A majority of the differentially expressed PCGs at each one of the three time points were also detected in at least one another time point. **(C)** Differentially expressed lncRNAs and Venn diagram of differentially expressed lncRNAs at three stages. **(D)** Hierarchical clustering heatmap and dendrogram of both mT/mG tumor and mT/mG control groups.

### 2.8 Correlation Analysis Between lncRNA and PCGs

The *cis*-regulatory potential target PCGs by lncRNAs were identified by the following procedures. For each lncRNAs, we identified PCGs as “*cis*-regulated PCGs” when: (1) the mRNAs locus was within 500k windows up- or downstream of the given lncRNA, and (2) the Pearson correlation of lncRNA-PCGs expression was significant (P-value of correlation less than 0.01). For each pair, Pearson correlation was performed to assess the correlation. Pearson correlation coefficients were calculated using cor() function and P values were obtained by corr.test() from library(‘psych’) in R.

### 2.9 *LINC00536*/*TRPS1* Expression Analysis

Gene expression profiles of *LINC00536*/*TRPS1* were analyzed by the web server, GEPIA2 (gene expression profiling and interactive analyses, version two; http://gepia2.cancer-pku.cn/~index), with the following parameters: |Log2FC| Cutoff: 1, P-value Cutoff: 0.01, log scale: log2(TPM + 1) matching TCGA normal and GTEx data ^29^.

### 2.10 Survival Curve Analysis

We utilized the “Survival Analysis” module of GEPIA2 to perform the overall survival curve analysis of *LINC00536* and *TRPS1* using both the TCGA-PRAD and GTEx BC datasets, respectively. The group cutoff of “Median” and axis units of “Months” were used. The plots with 95% confidence interval, P value of logrank test, HR (hazards ratio), and P value of Mantel-Cox test were generated ^29^.

### 2.11 External Validation in scRNAseq Dataset

Annotated cell types of the GEO: GSE176078 datasets as well as their clinical information were all obtained from their previous study ^30^. “Seurat” R package (v5) was applied to accomplish subsequent analysis ^31^. We selected the top 2000 highly variable genes (HVGs). The process involved the utilization of the top 10 principal components, in conjunction with HVGs. We used UMAP to reduce the dimensions and observe the classification of each cell type. According to previously reported epithelium markers, cancer epithelial cells and normal epithelial cells were further distinguished ^30^.

### 2.12 Cell Culture

Human MCF-7 (cat#HTB-22) and SK-BR-3 (cat#HTB-30) BC cell lines acquired from the American Type Culture Collection (ATCC, Manassas, VA, USA) were sequenced-validated. MCF-7 was cultured in RPMI1640 medium (cat#350-006CL, Wisent Inc., St. Bruno, QC, Canada) supplemented with 15% FBS (cat#080150), 2mM L-Glutamine (cat#609-065-EL) + 0.01mg/ml human recombinant Insulin (cat#521-016-IL), 100 IU/mL penicillin, and 100 ug/mL streptomycin (cat#450-201-EL). SK-BR-3 was cultured in McCoy’s 5A Medium (cat#30-2007, ATCC) supplemented with 10% FBS (cat#080150), 2mM L-Glutamine (cat#609-065-EL), 100 IU/mL penicillin, and 100 ug/mL streptomycin (cat#450-201-EL). Cells were tested and confirmed negative for mycoplasma every 6 months using the ZmTech Mycoplasma PCR/RT-PCR Detection Kits (Cat. M208001/M208002). The cells were not passaged more than 20 times; new vials were thawed.

### 2.13 Quantitative Real-Time PCR (qRT-PCR) Assays

Single-stranded cDNA was synthesized from 1 µg of total RNA using 5X All-In One RT MasterMix Kit (Abm, Vancouver, BC) followed by qPCR with PowerUp SYBR Green Master Mix (Applied Biosystems, Foster City, CA, USA) on a ViiA 7 Real-Time PCR System (Applied Biosystems). A standard program for the qPCR thermocycler was performed according to the previously published detailed protocol^19^. Human HPRT were used as endogenous controls to normalize each sample. The samples were run in triplicate. Primer sequences are provided in Supplementary Table S3.

### 2.14 Targeted SiRNA Knockdown

Human BC cell lines were transfected with Silencer® Select siRNA against *LINC00536* (Assay ID: N546107, Thermo Fisher Scientific) 24 h after plating. After the former titration, we applied 10 nM of *LINC00536* siRNA by use 2.5 µl per well (12-well plate) Lipofectamine RNAiMAX Transfection (Catalog#: 13778030, Thermo Fisher Scientific) following the manufacturer’s instructions (Thermo Fisher Scientific). A siRNA with a scrambled sequence (Catalog#: 4390843, Thermo Fisher Scientific) was used as nonsense control at the same concentration as the targeting siRNA. The plates were incubated in a 5% CO2 incubator at 37 ^◦^C for 48h and RNA was isolated using the RNeasy 96 kit (QIAGEN) according to the manufacturer’s instructions. qRT-PCR was performed to check the knockdown efficiency.

### Data Availability Statement

The datasets generated and analyzed for RNASeq in this publication, corresponding raw read counts matrix, and normalized gene expression matrix reported in this study have been deposited in NCBI’s Gene Expression Omnibus (GEO) ^48^ and are accessible through GEO Series accession number GSE225877. The published dataset accession number (GSE225877) corresponds to the data generated in the present study.

Publicly available data generated by others were used by the authors: the data analyzed in this study were obtained from Gene Expression Omnibus (GEO) at GSE176078, GSE76772, TCGA-PRAD and GTEx BC. All other raw data generated in this study are available upon request from the corresponding author.

## 3. Results

### 3.1. Generation of PyMT BC Mouse Model and Enrichment of Mammary Epithelial Cells

First, we generated MMTV-Cre; mT/mG mice by crossing the MMTV-Cre and mT/mG mice (Figure 1A). We termed MMTV-Cre; mT/mG mice as mT/mG control in which the mammary epithelium, expressing membrane-targeted GFP, is distinguished against the membrane-targeted red fluorescent backlight of stromal and nonepithelial-derived mammary gland tissues. Next, we combined the most widely used BC mouse model, namely MMTV-PyMT, with mT/mG control mice, which allowed us to specifically trace and enrich the GFP^+^ mammary epithelial cells from the early stage of tumor development to late-stage carcinoma. We termed MMTV-Cre; mT/mG; MMTV-PyMT as mT/mG tumor. These resulted in the generation of mT/mG, PyMT-induced breast tumors and their non-cancer controls (Figure 1A). From our previously published manuscript, H&E staining of mT/mG tumor mice displayed no histological difference at different stages of tumor progression compared with the original MMTV-Cre tumor mice ^17^. To focus our attention on changes in gene expression exclusively in the epithelium, we first isolated tumor tissues from mT/mG tumor mice as well as the normal mammary gland tissues from mT/mG control mice. We then processed these tissues to single cells *in vitro* and FACS-sorted them based on GFP expression (Figure 1B). Given the expression of GFP in both the mT/mG tumor and non-cancer mice, we were able to compare the transcriptomes of GFP^+^ BC cells at 6-, 8-, and 10-weeks and GFP^+^ normal mammary epithelial cells at the same time points. We also stained the intact tissues with the epithelial marker CK8/18 (green) and the basal/myoepithelial marker CK5 (red) in figure 1C which clearly shows intact structure of the mammary epithelium at an early stage of tumor progression. In this model, Cre is specifically expressed in the mammary epithelium^6,17^ resulting in GFP expression in mT/mG mice. It should also be noted that a small number of cells are double positive (expressing both GFP and mTd). This may have been the result of cells in which the expressed mTd protein after Cre excision has not yet been completely removed or from recombination occurring in only one mTmG allele (but not both) in homozygous mice.

### 3.2. DEGs in BC Initiation and Progression

We collected both GFP^+^ epithelial and GFP^+^ BC cells at weeks 6, 8 and 10 for bulk RNAseq analysis. We first performed PCA analysis (see Methods). The GFP^+^ tumor samples are distinctly separated from the GFP^+^ normal mammary epithelium samples on PC1. There is also a temporal separation among the GFP^+^ tumor samples at the 3 stages except for one tumor sample at week 10 on PC2 (Figure 2A). Next, we examined DEGs at each stage. At each time point (week 6, 8, and 10), we compared three mT/mG tumor samples with two mT/mG controls, and identified 1,120, 1,670, and 1,045 DEGs respectively (fold change > 2, false discovery rate (FDR) < 0.1). Among these, 992 mRNAs (276 up-regulated and 716 down-regulated) were differentially expressed at week 6; 1,440 mRNAs (306 up-regulated and 1,134 down-regulated) were differentially expressed at week 8; and 926 mRNAs (304 up-regulated and 622 down-regulated) were differentially expressed at week 10 (Figure 2B). RNAseq results also identified that 128 lncRNAs (67 up-regulated and 61 down-regulated) were differentially expressed at week 6; 230 lncRNAs (83 up-regulated and 147 down-regulated) were differentially expressed at week 8; and 119 lncRNAs (62 up-regulated and 57 down-regulated) were differentially expressed at week 10 (Figure 2C). Among the union of all 2,237 DEGs, a significant proportion of DEGs (24.7%, 472 PCGs, and 13.0%, 42 lncRNA genes) appeared at all three stages (Figure 2B & 2C).

Our methodology which involves *in vitro* mammary epithelial cell growth before enrichment and isolation of GFP+ cells by FACS for RNAseq may potentially induce signaling pathway changes which may affect data interpretation. In order to examine this possibility, we compared our results with a published RNAseq dataset (GSE76772), which used intact mammary tumors and normal gland tissues from PyMT mice and age-matched controls at three time points similar to our study i.e. hyperplasia at week 6, adenoma at week 8, and carcinoma at week 10 ^32^. First, our PCA analyses showed that the tumor cell samples were clearly clustered and separated from the normal epithelial cell samples, which is in agreement with the GSE76772 dataset (Figure 2A) ^32^. Secondly, we also observed that more genes were downregulated than upregulated in the tumor cell samples at each time point and a significant number of DEGs overlapped at three time points analysed in our dataset (Figure 2B & 2C) ^32^. Thirdly, our comparative RNA-seq analyses revealed that GFP^+^ cells after 48h of *in vitro* culture closely resembles the gene expression signature of the intact tissues derived from this same mouse model ^32^. All the breast cancer related genes are significantly highly expressed in tumour samples compared to non-tumour samples (Figure 2D). For example, genes regulating cell cycle and proliferation, *Akt1*, *Ccnd1*, *Tacc3*, *Mcm6*, and *Mki67* and genes playing a crucial role of DNA methylation in tumor development were significantly overexpressed in our tumor compared to control mice (Figure 2D).

### 3.3. Time-Course Analysis of mRNAs and lncRNAs Expression Patterns in BC Initiation and Progression

To investigate the temporal expression patterns of mRNAs and lncRNAs during breast tumor initiation and progression, the union of 324 differential expressed lncRNAs and 1,913 differentially expressed mRNAs were used as input data to analyze patterns in gene expression. Based on the sum of squared errors (SSE), the DEGs were divided into four clusters: cluster 1 included 866 genes, cluster 2 had 191 genes, cluster 3 had 496 genes, and cluster 4 had 682 genes (Fig. 3A). Genes in cluster 1 were dramatically reduced expression in the hyperplasia stage (week 6) and further maintained a low expression as the tumor progressed (week 8 to 10). Genes in cluster 2 showed significantly increased expression especially at hyperplasia stage (week 6) but gradually decreased as the tumor progressed (week 8 to 10). Meanwhile, genes in cluster 3 increased expression gradually and remained at a high level at weeks 8 and 10; whereas genes in cluster 4 were reduced expression at hyperplasia and adenoma stages (weeks 6 and 8) and then increased to base-line level at carcinoma stage (week 10). A line graph showing kinetics of changes in gene expression profile within a given cluster at normal, hyperplasia, adenoma, and carcinoma stages, is shown in Figure 3.

**Figure 3.**
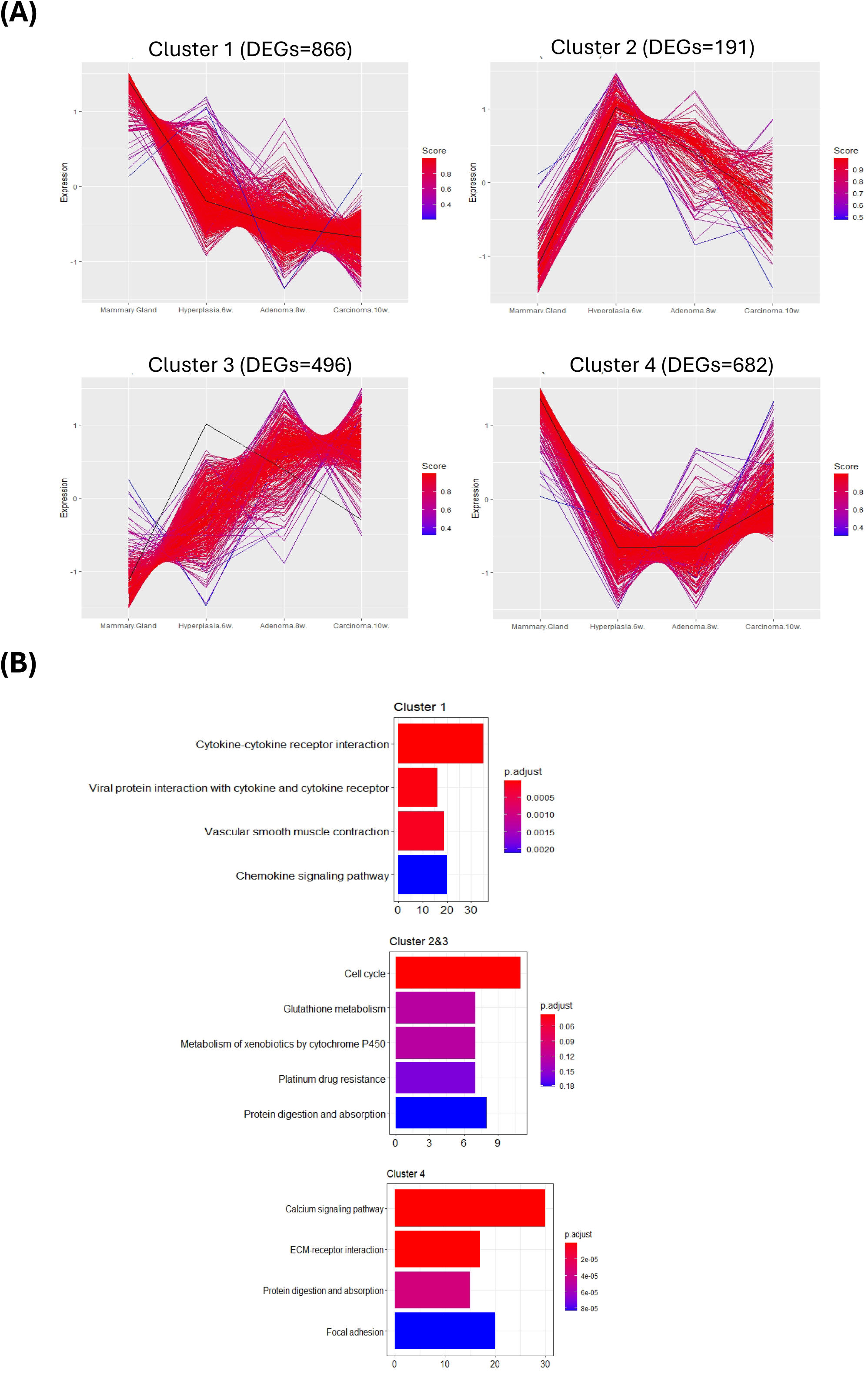
K-means clustering analysis of differentially expressed mRNAs and lncRNA genes from normal, hyperplasia, adenoma, and carcinoma stages of tumor progression. **(A)** Line graphs showing kinetics of changes in gene expression profile within a given cluster (K = 4). Each line indicates one DEG. The red line indicates the high similar one and the blue line indicated the low similar one. **(B)** KEGG enrichment of the four K-means clusters. The top 5 results are plotted.

To further understand the potential functional role of DEGs with obvious temporal correlation, gene enrichment analysis of 4 clusters was performed. As shown in Figure 3B, cytokine-cytokine receptor interaction (KEGG: mmu04060) was enriched in cluster 1. Cell cycle (KEGG: mmu04110) and glutathione metabolism (KEGG: mmu00480) were enriched in clusters 2 and 3. Genes in cluster 4 played roles in the calcium signaling pathway (KEGG: mmu04020), the ECM-receptor interaction (KEGG: mmu04512) and during protein digestion and absorption (KEGG: mmu04974). Collectively, these results suggest that the genes have similar expression patterns (co-expression genes) which may play a similar functional role through the same regulatory mechanisms (co-regulated genes) in BC initiation and progression.

### 3.4. Characterization of Aberrantly Expressed lncRNAs in BC Initiation and Progression

We identified a total of 324 lncRNAs differentially expressed in BC cells derived from mT/mG tumor mice compared to normal mammary epithelial cells from mT/mG control mice at 3 different stages (Figure 2C). These lncRNAs represent a diverse group in terms of their biotypes and genomic location (Figures S1A). Biotypes include ‘‘lincRNA’’ (long intergenic non-coding RNAs), ‘‘processed pseudogenes’’, ‘‘antisense’’ transcripts, “TEC” (to be experimentally confirmed), and ‘‘processed transcripts’’ (Figure S1A). We performed coding potential analysis on all 324 differentially expressed lncRNAs using the Coding-Potential Assessment Tool (CPAT) ^33^. Our results indicated that 84% of the 324 identified transcripts do not exhibit coding potential despite including pseudogenes in our analysis (Figures S1B). A complete list of their official gene IDs, gene name, biotypes and coding potential are presented in Table S1. Interestingly, very few of the identified mouse lncRNA transcripts have been studied previously. Among the identified ones, *C130071C03Rik*, *A730020E08Rik* and *Foxd2os* have shown at least 2-fold upregulation in PyMT-induced organoid tumors compared to organoid from normal mammary epithelium ^34^.

To explore the potential *cis*-regulatory functions of lncRNAs in BC, we performed Pearson correlation coefficient analysis to evaluate the correlation between lncRNAs and PCGs, and the lncRNAs and PCGs with Pearson correlation coefficient no less than 0.80 and the P value less than 0.01 were further mapped to their genomic locus. LncRNAs which locate within 500K ranges upstream or downstream of the PCGs were screened further. A total of 93 lncRNA-PCGs pairs were obtained from the lncRNA-PCGs co-expression analysis and are shown in Table S2. Among them, 91 lncRNA–PCGs pairs showed positive correlations, with only 2 pairs showing negative correlation. To further confirm the relevance of mouse lncRNAs in human BC, we started to identify human orthologs of the 93 lncRNA-coding gene pairs. We compared mouse and human transcripts at the level of both sequence conservation and genomic location. If the mouse lncRNA sequence matched an annotated human gene using BLAT alignment tool, then the respective transcript was regarded as the human counterpart. We also compared the neighboring genes of each lncRNA and included their human counterpart based on genomic location rather than on sequence considering their functional conservation ^35^. We matched 23 potential annotated human lncRNAs from our 93 mouse pairs (Table 1). Among the 23 annotated human lncRNA-PCG pairs: 11 pairs came from cluster 1; 7 from cluster 3; and 5 from cluster 4 (Figure 3A).

**Table 1.** Human counterparts of mouse lncRNAs.

| LncRNAs |  |  | Protein Coding Genes |  |
| --- | --- | --- | --- | --- |
| Mouse gene Name | Human counterpart (seq, hg38) | Human counterpart (synteny, hg38) | Gene Name | Description |
| **Cluster 1** |  |  |  |  |
| *AI504432* | *ENSG00000288847* | *ENSG00000288847* | *KCA3* | Potassium voltage-gated channel subfamily A member 3 |
| *C130021I20Rik* | *RP11-123K19.1* | *RP11-123K19.1* | *LMX1B* | LIM homeobox transcription factor 1 beta |
| *Dio3os* | *DIO3OS* | *DIO3OS* | *DIO3* | Iodothyronine deiodinase 3 |
| *Fzd10os* | *FZD10-AS1* | *FZD10-AS1* | *FZD10* | Frizzled Class Receptor 10 |
| *Gm2670* |  | *WNT5A-AS1* | *WNT5A* | Wnt family member 5A |
| *Gm41609* |  | *DLGAP1-AS1* | *DLGAP1* | Disks large-associated protein 1 |
| *Gm42481* | *ENSG00000249678* | *ENSG00000249678* | *PCDH7* | protocadherin 7 |
| *Gm42679* |  | *NGF-AS1* | *NGF* | Nerve growth factor |
| *Gm9913* | *FBN1-DT* | *FBN1-DT* | *FBN1* | Fibrillin 1 |
| *Hoxb3os* | *HOXB-AS1* | *HOXB-AS1* | *HOXB3* | Homeobox protein Hox-B3 |
| *Tbx3os1* |  | *TBX3-AS1* | *TBX3* | T-box transcription factor 3 |
| **Cluster 3** |  |  |  |  |
| *BC016548* |  | *LINC02707* | *ELF5* | E74 like ETS transcription factor 5 |
| *Gm12860* |  | *ENSG00000286668* | *CITED4* | Cbp/p300 interacting transactivator with Glu/Asp rich carboxy-terminal domain 4 |
| *Gm19303* | *LINC00536* | *LINC00536* | *TRPS1* | Transcriptional Repressor GATA Binding 1 |
| *Gm26814* |  | *PPP1R9A-AS1* | *PPP1R9A* | Protein phosphatase 1 regulatory subunit 9A |
| *Gm50337* |  | *ENSG00000231188* | *SCD1* | Stearoyl-CoA desaturase |
| *Pinc* | *DIRC3-AS1* | *DIRC3-AS1* | *TNS1* | Tensin 1 |
| *Ttc39aos1* |  | *TTC39A-AS1* | *TTC39A* | Tetratricopeptide repeat domain 39A |
| **Cluster 4** |  |  |  |  |
| *AC109619.1* |  | *RP11-21A7A.3* | *FLRT1* | Fibronectin leucine rich transmembrane protein 1 |
| *Gm13631* |  | *MYO3B-AS1* | *MYO3B* | Myosin IIIB |
| *Gm14377* | *RP11-1081L13.6* | *RP11-1081L13.6* | *PTPN5* | Protein tyrosine phosphatase non-receptor type 5 |
| *Gm17396* | *RBMS3-AS3* | *RBMS3-AS3* | *RBMS3* | RNA binding motif single stranded interacting protein 3 |
| *Gm49417* | *RP11-582J16.4* | *RP11-582J16.4* | *SORBS3* | Sorbin and SH3 domain containing 3 |

### 3.5. Assessment of Clinical Relevance of Human *LINC00536*/*TRPS1* in BC

Here, we focus on *Gm19303*, as this mouse lncRNA in cluster 3 gradually increased its expression as the tumor progressed. BLAT results for *Gm19303* indicate sequence identity in the human genome as well as a similar genomic context. It matches a longer lncRNA, *LINC00536*, in close spatial proximity of two PCGs, upstream of *TRPS1* and downstream of *EIF3H* (Figure 4A). *LINC00536* contains 14 exons, and its sequence spans 2,692 bp in the human genome with a sequence identity of 92.4% and a BLAT score of 422. This genomic region is highly conserved throughout vertebrates ^36^. Pearson correlation analysis indicate *Gm19303* has a specific correlation with neighboring PCGs, Trps1 (Pearson correlation coefficient r = 0.93, P value = 7.21E-06), indicating that *Gm19303* has a potential *cis*-regulatory effect on *Trps1* (Table S2).

**Figure 4.**
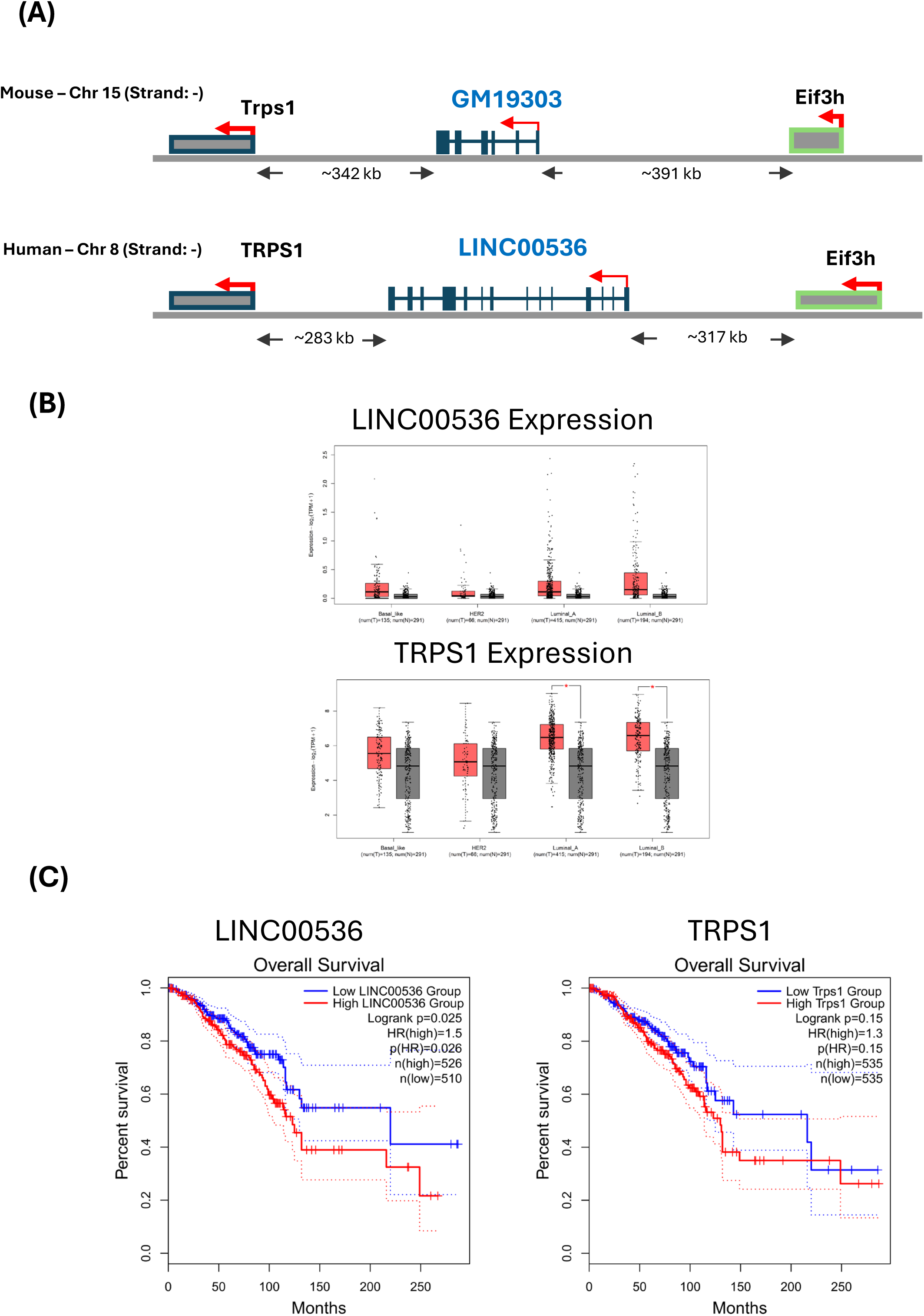
*LINC00536* and *TRPS1* are overexpressed in BC. **(A)** Identification of a human *LINC00536* gene based on sequence identity and genomic location. The upper panel represents the mouse genome (mm10); the lower panel represents the human genome (hg38). *LINC00536* is an intergenic lncRNA gene located on human chromosome 8. **(B)** Corresponding box plots of the comparative expression of *LINC00536* (upper panel) and *TRPS1* (lower panel) in BC samples (red) versus normal tissue samples (grey) generated using GEPIA2 in 4 subgroups of BC. **(C)** BC survival analysis plots (Kaplan-Meier) were generated for *LINC00536* (left panel) and *TRPS1* (right panel) using GEPIA2.

To evaluate the expression status of the human lncRNAs, *LINC00536*, we applied GEPIA2 ^29^ to analyze RNAseq data of The Cancer Genome Atlas (TCGA) and The Genotype-Tissue Expression (GTEx) databases by comparing BC samples to matched normal tissue samples. We found both *LINC00536* and *TRPS1* to be significantly overexpressed in breast tumors compared to normal breast tissue (Figure 4B), confirming our initial RNAseq screen in the PyMT-induced BC mouse models. Interestingly, both *LINC00536* and *TRPS1* are exclusively expressed in breast tumor samples but not in other types of cancers, indicating that *LINC00536* and *TRPS1* expression appears to be lineage-restricted (Figure S2). We further investigated whether *LINC00536*/*TRPS1* are expressed in a subtype-specific manner by analyzing all TCGA BC samples with subtype information. We observed statistically significant differences in the expression level of *TRPS1* comparing different subtypes of BC, with highest levels in the luminal A and B subtypes, and lowest levels in the HER2^+^ subtype (Figure 4B). Survival analysis indicated that patients with high levels of *LINC00536* are correlated with poorer BC patient survival outcomes, perhaps indicating an oncogenic role (Figure 4C). Following a similar pattern, high expression of *TRPS1* is associated with slightly poorer BC survival from TCGA and GTEx databases (Figure 4C). Neither *LINC00536* nor *TRPS1* expression is associated with patient survival for other types of cancers (Figure S3).

### 3.6. Assessment of Human *TRPS1* Expression in Atlas Human BCs

We found that *LINC00536*/*TRPS1* were highly expressed in BC tumor cells, and this was associated with poor prognosis, suggesting a potential diagnostic value. However, there is limited research on their expressions in BC at the single cell level. Therefore, the scRNAseq data (GSE176078) was utilized to validate our current findings. The GSE176078 was a scRNAseq dataset included a total of 26 BC samples, containing 11 ER^+^, 5 HER2^+^, and 10 TNBCs. First, we performed analysis of scRNAseq data to further clarify the localization of *LINC00536*/*TRPS1* and their expression pattern at the single cell level. The cells were annotated using canonical lineage markers: endothelial cells, epithelial cells, and other immune cells (Figure 5A). By locating the *TRPS1* expression, it was primarily expressed in epithelial cells (EPCAM; Figure 5B). A total of 8,917 epithelial cells were re-clustered, including 1,960 normal epithelial cells and 6,957 cancer epithelial cells (Figure 5C). *TRPS1* was highly expressed in cancer compared to normal epithelial cells (Figure 5D). We re-clustered 6,957 cancer epithelial cells from the 11 ER^+^, 5 HER2^+^, and 10 TNBCs samples (Figure 5E). We found that *TRPS1* were highly expressed in ER^+^ subgroup compared to HER2^+^ or TNBC subgroups (Figure 5F). Our scRNAseq analysis further confirmed the expression pattern of *TRPS1* in BC. However, unlike PCGs, lncRNA are expressed at a low level, and therefore, cell-specific *LINC00536* transcripts are hard to distinguish from technical noise at a single cell level due to inherent limitations such as dropouts of current scRNAseq techniques ^37^.

**Figure 5.**
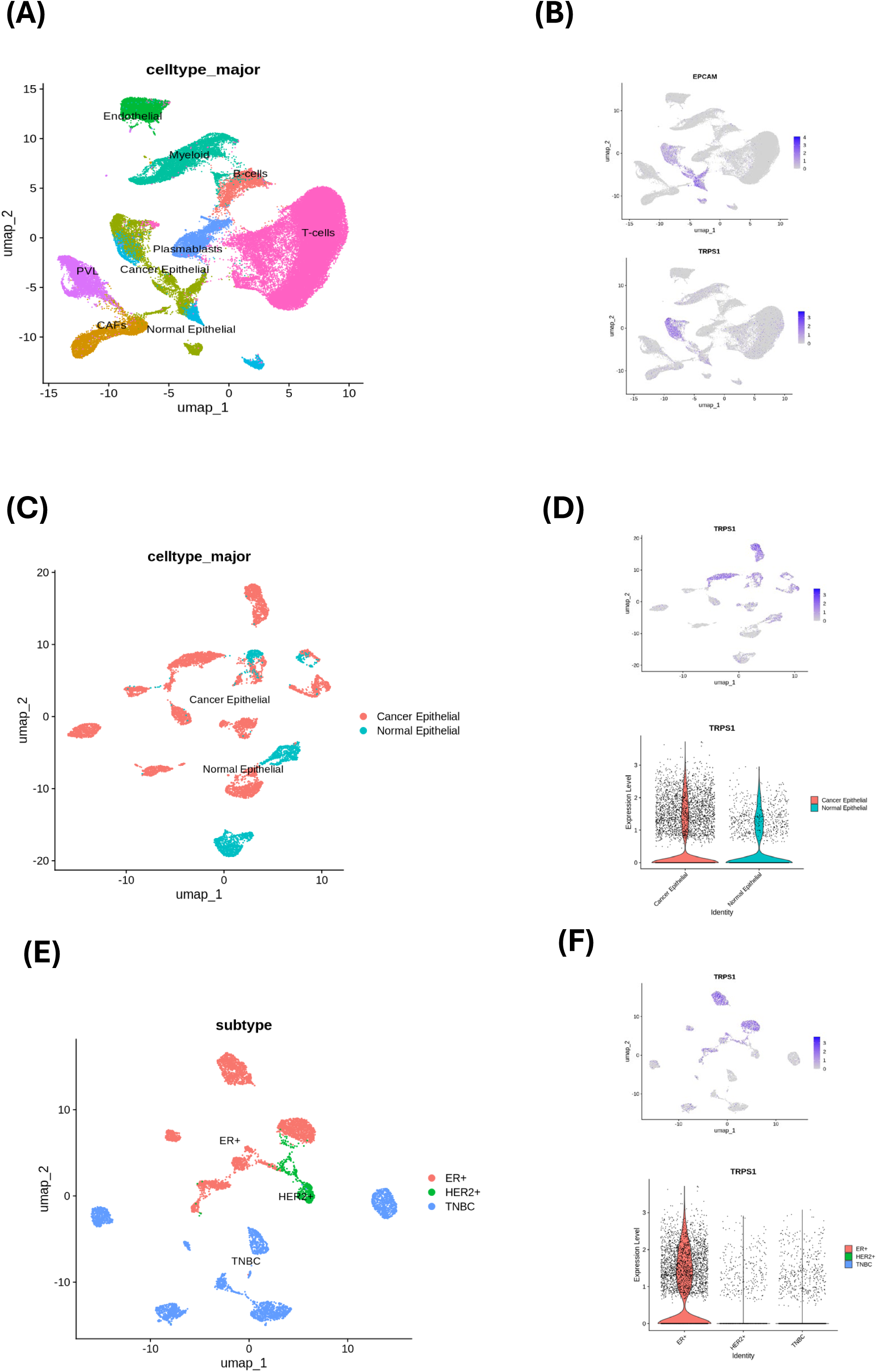
Cellular composition of primary BCs and *TRPS1* expression. **(A)** UMAP visualization of 67,588 cells analyzed by scRNAseq and integrated across 26 primary BCs. Clusters were annotated for their cell types as predicted using canonical markers. **(B)** Log-normalized expression of markers for epithelial cells (EPCAM) and *TRPS1*. **(C)** UMAP visualization of 8,917 epithelial cells colored by cancer or normal cells. **(D)** Log-normalized expression of *TRPS1* in all epithelial cells. **(E)** UMAP visualization of 6,957 cancer epithelial cells colored by clinical subtypes. **(F)** Log-normalized expression of *TRPS1* in all cancer epithelial cells.

### 3.7. Interruption of *LINC00536-TRPS1* Decreases Cell Proliferation Capacity

To validate the relevant role of the *LINC00536-TRPS1* in Human BC development, we next performed *in vitro* functional studies by knocking down this novel *LINC00536* in human MCF-7 and SK-BR3 BC cell lines using targeted siRNA. Our mouse RNAseq analyses suggested that *Gm19303* can potentially cis-regulate nearby *Trps1* and might contribute to BC progression. *LINC00536* gene is located on human chromosome 8 (strand-) adjacent to *TRPS1* (Figure 6A). We tested two different specific siRNAs and achieved knockdown efficiencies of *LINC00536* of almost 50% (48h) and more than 70% (72h) for both human MCF-7 and SK-BR3 cell lines using 10 nM of the most potent siRNA (Figure 6B&C). Concomitantly, we observed more than 45% decrease in *TRPS1* expression at both 48 h and 72 h for MCF-7 cell line (Figure 6B), indicating that *LINC00536* can cis-regulate *TRPS1* expression.

**Figure 6.**
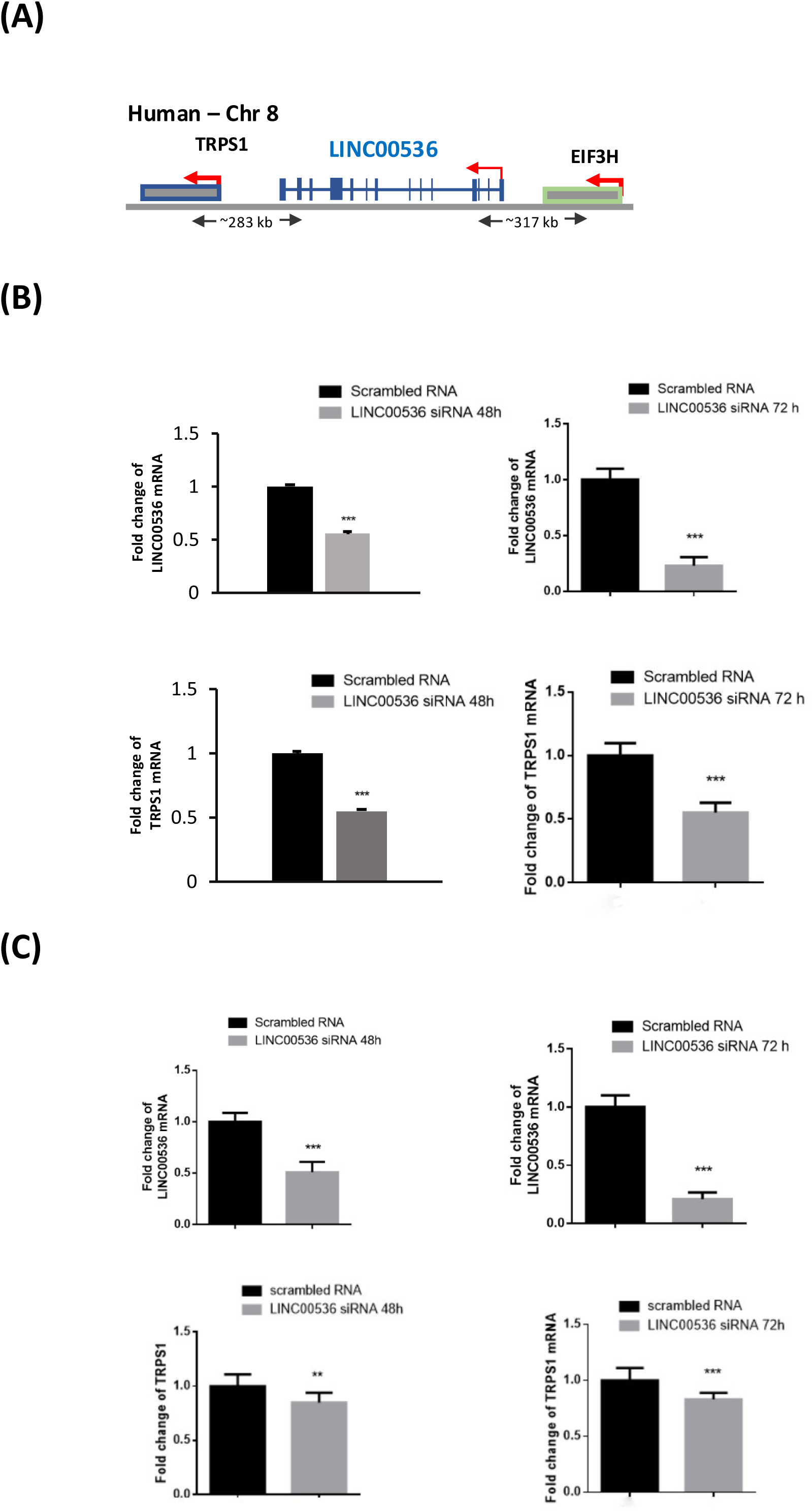
Specific siRNA knockdown of *LINC00536*. (**A**) Representation of the *LINC00536* gene locus. *LINC00536* is an intergenic lncRNA gene located on Human chromosome 8, and the *LINC00536* RNA transcript contains 14 exons and a poly (**A**) tail. (**B**) qPCR showing *LINC00536* and *TRPS1* mRNA expression in human MCF-7 BC cell line after 48 h and 72 h of incubation with *LINC00536*-specific siRNA compared to scrambled siRNA-treated control cells. Data are presented as mean values ± SD (*n* = 3 independent experiments). * *p* < 0.05 (paired student’s *t*-test; two tailed.) (**C**) qPCR showing *LINC00536* and *TRPS1* mRNA expression in human SK-BR3 BC cell line after 48 h and 72 h of incubation with *LINC00536*-specific siRNA compared to scrambled siRNA-treated control cells. Data are presented as mean values ± SD (*n* = 3 independent experiments). * *p* < 0.05 (paired student’s *t*-test; two tailed.)

Additionally, knockdown of *LINC00536* by targeted siRNA significantly decreased TRPS1 protein levels compared to the control group for both the human MCF-7 and SK-BR3 BC cell lines, validating the downstream impact of *LINC00536* knockdown (Figure S4A). To further investigate the functional impact of *LINC00536* downregulation on tumor cell proliferation, knocking down *LINC00536* reduced Ki67 expression both in human MCF-7 and SK-BR3 BC cell lines (Figure S4B). Our *in vitro* results therefore validated the relevance of *LINC00536-TRPS1* overexpression to tumor cell growth.

## 4. Discussion

In the present study, we generated a PyMT-induced BC mouse model in which mammary epithelial cells are specifically traced by membrane-targeted GFP (mT/mG reporter). This BC mouse model allowed us to specifically enrich mammary epithelial cells, including both normal and pre-neoplastic cells. Therefore, we were able to perform an unbiased RNAseq screen to identify all the differentially expressed lncRNAs in mammary tumor compared to normal mammary epithelial cells during BC initiation and progression. In our screen, we identified 324 differentially expressed lncRNAs, most of which have not been described previously in the context of BC. Our study emphasizes the importance of unbiased genome-wide gene expression approach and unravels valuable candidates lncRNAs as potential prognostic and/or therapeutic targets in mammary carcinogenesis. Our RNAseq screen identified many proteins coding DEGs in mammary tumor cells compared to normal controls. The integration analysis of lncRNA-PCGs is one of the most common approaches to predict lncRNAs functions in different biological processes.

Therefore, we applied the correlation analysis of the differentially expressed lncRNAs and PCGs and identified a total of 93 lncRNA-PCGs pairs which are abundantly correlated between the transcripts of the two groups. Based on the relationship between lncRNAs and their associated PCGs, lncRNAs can be further classified into several categories, including intergenic, intronic, bidirectional, and sense and anti-sense ^38^. Previous studies have shown that lncRNA can mainly act in both *cis* and *trans* to regulate the expression levels of target genes, which is the major regulatory pattern of lncRNA in higher organisms ^10^. Many lncRNAs regulate nearby PCGs expression through epigenetic, transcriptional, or post- transcriptional mechanisms ^39^. Comprehensive analysis of lncRNAs and PCGs profiles provides us with a better understanding of biological functions of these differentially expressed lncRNAs.

We infer that the human counterparts of lncRNAs which we identified from our mouse model likely impact human BC initiation and progression. Our lncRNA PCG profile revealed concomitant up-regulation of *Gm19303* and *Trps1* in mouse BC. To evaluate the expression status of their potential human counterparts, *LINC00536* and *TRPS1*, we analyzed RNAseq data of TCGA and GTEx datasets by comparing BC to matched normal tissues. Notably, *LINC00536* and *TRPS1* were found to be upregulated in BC, confirming our initial RNAseq screen in our mouse model. Additionally, *LINC00536* and *TRPS1* showed strong luminal subtype specificity, indicating that they may be clinically relevant in certain subtypes.

In a recent case-control GWAS study, a total of 13 statistically significant subtype-informative risk loci has been identified. Among them, *8q23.3*/*LINC00536*/*TRPS1* is one of fine mapping implicated functional/causal variants and risk genes for TNBC as compared with luminal cancer ^40^. At *8q23.3*, nine variants are located within or directly adjacent to *LINC00536*, five of which are reside in the active enhancer regions in BC cell lines or human mammary epithelial cells ^40^. One of the variants, *rs2223054*, maps to a ∼5 kb genomic segment which can interact with the *TRPS1* promoter region ^40^. *TRPS1* is a transcriptional repressor which plays an indispensable role for luminal epithelial cell survival in mammary gland development as well as involved in BC development ^41^. *LINC00536* has been shown to be overexpressed in BC and silencing this lncRNA inhibited cell proliferation, migration, and invasion capacities in human BC ^42^. The collected evidence demonstrated that *TRPS1* could be activated by *LINC00536* via long-range chromatin interactions at the *TRPS1* promoter region and such interactions between *TRPS1* and *LINC00536* could exert their biological functions in BC ^40^. We performed targeted siRNA knockdown of *LINC00536* and observed a significant reduction of nearby *TRPS1* expression, confirming that *LINC00536* has the potential to cis-regulate nearby PCGs in human BC cell lines. Our study using both computational approaches and molecular functional assays provides novel evidence that lncRNAs are important regulators during BC initiation and thereby play a critical role in BC progression. Future studies will be directed at elucidating the biological mechanisms underlying *LINC00536*-mediated transcriptional regulation in BC initiation and progression.

We identified total 23 annotated human lncRNAs by comparing mouse and human nucleotide sequence or synteny between non-coding and PCGs. Importantly, among them, several lncRNAs have been previously linked to BC. For example, lncRNA *TTC39A-AS1* is located at human chromosome 12 and transcribed in opposite directions from the *TTC39A* gene locus. *TTC39A-AS1* expression was overexpressed in BC than normal tissues based on TCGA database as well as in several BC cell lines ^43^. Moreover, patients with BC with a high level of *TTC39A-AS1* had a shorter overall survival than those with a low level of *TTC39A-AS1* ^43^. LncRNA *DIO3OS* is located at human chromosome 14 and transcribed in opposite directions from the *DIO3* gene locus. *DIO3OS* has been observed downregulated in almost all cancers, including liver and thyroid cancer ^44^. Furthermore, decreased *DIO3OS* expression tended to predict poor prognosis and lower survival rates in hepatocellular carcinoma ^45^. However, *DIO3OS* has been found overexpressed in aromatase-inhibitor-resistant BC, and its expression associated with a poor prognosis in these patients ^46^. Overexpression of *DIO3OS* in ER-positive BC upregulates lactate dehydrogenase A (*LDHA*) expression and promotes glycolytic metabolism by interacting with polypyrimidine tract binding protein 1 (*PTBP1*) and stabilizes the mRNA of *LDHA* ^46^. Our study suggests that multiple lncRNAs are aberrantly expressed in BC from our initial RNAseq screen in the PyMT-induced BC mouse models. However, the biological functions of these lncRNAs in human cancers warrant further investigations.

There were additional limitations in our study. First, previous genomic analysis indicated that PyMT-induced BC mouse model is initially associated with the mature luminal signature at an early stage then progresses to basal-like tumors with low levels of ER in later stages, which is associated with metastasis ^47^. In contrast, BC in women is characterized by significant heterogeneity of individual tumor tissues as well as different pathological and molecular subtypes that have different treatment responses and clinical outcomes.

Therefore, our results here cannot extend to all types of human breast cancer types. Moreover, our identification of human lncRNA counterparts, and their relevant functions also have some limitations. Many lncRNAs in mice are not highly conserved in humans which often hinders the identification of human orthologs via a mouse model. In our study, we included 23 annotated human lncRNAs counterparts by comparing human and mouse genome at the level of genetic location or sequencing conservation. Future studies are necessary to unambiguously validate the rest of this unannotated human counterpart from our 93 mouse lncRNA list. Thirdly, our study only focused on the lncRNAs with a *cis*-regulatory function, and future studies will be necessary to explore the lncRNAs with a potential *trans*-acting function. Finally, we only validated the overexpression of *LINC00536*/*TRPS1* in human BC, and future studies will be necessary to explore and validate the potential downregulated lncRNAs that we identified in our RNAseq screen from our mouse model which may exert a tumor suppressor function in human cancer.

## 5. Conclusion

We combined the classic PyMT-induced BC mouse model with MMTV-Cre; mT/mG, which allowed us to specifically trace and enrich the GFP^+^ mammary epithelial cells from the early stage of tumor development to late-stage carcinoma. We performed an RNAseq screen to examine the expression of lncRNAs in the full range of breast tumor development. We identified 324 lncRNAs that are aberrantly expressed at least 2-fold in mT/mG tumor samples. Among these transcripts we matched 23 pairs potential annotated human lncRNAs counterparts and their potential *cis*-regulatory target PCGs. We further confirmed that both *LINC00536* and *TRPS1* to be significantly overexpressed in human breast tumor compared to normal breast tissue. Overexpression of either *LINC00536* or *TRPS1* are associated with survival rate of BC patients.

## Acknowledgments

This research was funded by the United States Department of Defense (grant number W81XWH-15-1-0723) (R.K.), Canadian Institutes of Health Research (MOP-142287) (R.K.), the CFI project Canada’s Genome Enterprise (CGEn) 35444, 33408 and 40104 (J.R.) and Compute Canada Resource Allocation Competition (RAC) wst-164 (J.R.).

## Author Contributions

Conceptualization, R.Z., J.R. and R.K.; methodology, R.Z., D.B. and J.L.; software, R.Z. and D.B.; validation, R.Z., J.L. and D.B.; formal analysis, R.Z. and D.B.; resources, R.K. and J.R., writing—original draft, R.Z.; writing—review and editing, R.Z., J.R., and R.K. All authors have read and agreed to the published version of the manuscript.

## Institutional Review Board Statement

The use of mice in this project has been reviewed and approved by the McGill Facility Animal Care Committees (FACCs) (Protocol (AUP) # MUHC-7744).

## Conflicts of Interest

The authors declare no potential conflicts of interest.

## Notes

### Competing Interest Statement

The authors have declared no competing interest.

